# The impact of artificial linear features on the pollination of plant communities

**DOI:** 10.1101/2025.04.02.646872

**Authors:** Dongbo Li, Christopher F. Clements, Jane Memmott

**Author notes:** Current Address: School of Biology, University of Leeds, Leeds, United Kingdom (Dongbo Li).

## Abstract

Linear features, such as road verges, hedgerows can function as corridors for species dispersal. Previous studies showed that these features could enhance movement of pollinators such as bees, increasing pollination success of bee pollinated plants. However, it remains unclear if these effects extend to communities of plants pollinated by other pollinators.

We conducted a field experiment where we measured the pollination of phytometer plant communities linked by linear features. We constructed six 30m artificial linear features in both urban and rural meadows in the Southwest UK, and placed plant communities on either end of these linear features, comparing these to phytometers without linking linear features. Plant communities were assembled with seven species known to attract different groups of insect pollinators including bees, flies, and moths. We tested self-incompatibility of plant species and used their seed set to measure pollination.

We found that linear features significantly improved the pollination success of three bee pollinated species, whilst the seed production of fly and moth pollinated species were either unaffected or less affected. Overall, it is likely that linear features benefit the pollination of some plant species more than others.

Our results suggest that linear features play an important role in maintaining pollination for some plant species, highlighting the importance of evaluating community-level impacts of corridors on pollination. Given the importance of bees as pollinators, linear features can provide a practical strategy to conserve and improve pollination, but where species are fly or moth pollinated, alternative strategies may need to be considered.

## 1. Introduction

The loss and fragmentation of natural habitats are one of the commonest stressors in human dominated ecosystems [1]. Habitat destruction and fragmentation can cause biodiversity loss, with ensuing loss of ecological functions [2]. One solution to mitigate the usually negative effect of fragmentation is to connect isolated patches with corridors, which can reduce the likelihood of extinction by promoting dispersal and increasing gene flow. The effectiveness of corridors depends on both their physical properties such as length, width, and quality [3–5] and on the ecology and behaviour of the species using the corridors [6]. Species may not respond to corridors equally as their response depends on factors such as their habitat preferences and dispersal behaviour [7, 8]. Given this, it is important to understand how different species respond to corridors, if we are to understand their impact.

Linear features, such as road verges, hedgerows, and forest edges can function as corridors for a diverse range of animals, including bees [9–11], moths [12], and small mammals [13, 14]. These linear features not only provide habitats and/or resources used for dispersal, but also serve as movement conduits which may direct their movement routes. For example, Brebner, Makinson [15] found that bumblebee *Bombus terrestris* used paths as landmarks to target resource search and Cranmer, McCollin [11] found that artificial hedgerows (i.e., no resources provided) could facilitate the movement of bumblebees (*Bombus spp.*). These types of features may play an important role in maintaining the pollination of bee-pollinated plants, as the movement of bees along linear features could increase their fitness by promoting pollen dispersal [9, 16]. While there is some information on this mostly on bees, our understanding of the importance of linear features on other important groups pollinators such as flies and moths remains very limited. Given the importance of flies and moths as pollinators of both crops and wild plants [17–19], understanding how these groups respond to linear features is important.

Insect pollinators consist of a range of species with fundamentally different ecologies. For instance, bees are central place foragers and need to return to their nests after foraging, which requires an integration of cognitive abilities such as visual learning, memory, and navigation [20]. In contrast, flies, moths, and butterflies are not central place foragers and tend to be dispersive, spending most of time searching food resources and finding sites for reproduction [21, 22]. However, moths can use linear structures to direct their movements [12], and some butterflies and flies respond to habitat boundaries as dispersal corridors [23–25], although this effect depends on the quality of surrounding habitats and species behaviour [6, 26]. Overall, different behaviours and ecologies within insect communities are likely to lead to different responses of pollinators to the same landscape structure [27], presenting a challenge for conserving these pollinators and the important pollination services they provide.

Most research on the effectiveness of biological corridors to plants and pollinators has focused on individual plant species and there are very few studies which measure the effect of corridors at the plant community level. This may overlook corridor effects on pollination due to the varying response of different types of pollinators to landscape structure and features. In addition, as effective pollination at individual species can be modified by community factors such as plant competition for pollinators [28], quantifying impact of corridors on pollination needs data from plant communities as plants do not flower in isolation. Here we use a phytometer plant community consisting of a set of species with complementary pollinator traits provide a useful means to evaluate the community-level responses of pollinators to their environment. While phytometers have a long history in pollination biology for measuring pollination levels in the field, most studies were carried out using a single plant species [29–31], and there were few studies measuring pollination at community level [but see 32, 33].

Here we investigate the effect of linear features on the movement of insect pollinators and pollination services, using a community of phytometer plants consisting of complementary pollinator traits. We constructed replicate 30-meter linear features in open meadows in both urban and rural landscapes, and placed a replicate plant community containing seven plant species at each end of the linear feature. Three of our plant species were principally pollinated by bees, two by flies, and two by moths. Only one individual of each species was in each community (i.e. there was no cross pollination within an individual community), allowing us to measure pollination based on the movements of pollinators. Seed set in our plants linked by corridors was compared to control plant communities, where no linear feature was present, this experimental design enabling us to compare the effectiveness of corridors on the bee, fly and moth pollinated plants used in our study.

## 2. Materials and methods

### 2.1 Field sites

The experiment ran from July 2021 to September 2021 at two field sites, one urban and one rural, in the Southwest of England, where linear features are common landscape structures in both types of habitat [11, 34]. The urban site was located at the University of Bristol Student Residences (51°48’ N, 362° 62’ W) and the rural site was Fenswood Farm (51° 25’ N, 362° 40’ W), Bristol, United Kingdom. Both sites were meadows, ranging from 0.3 −5.2 ha, with the urban site surrounded by buildings and woodlands, and the rural one by mixed farmland (Figs. 1a & b). The urban meadows were regularly mown and the rural meadows were largely unmanaged, aside from a hay-cut in late July. The centre of the meadows consisted of low growing plants and most of the meadows had hedgerows around the edge. The centres were clear of all linear features. Insect pollinators included bumblebees (*Bombus sp.*), honey bees (*Apis sp*.), flies, beetles and butterflies were observed in both types of sites during the fieldwork.

**Fig 1.**
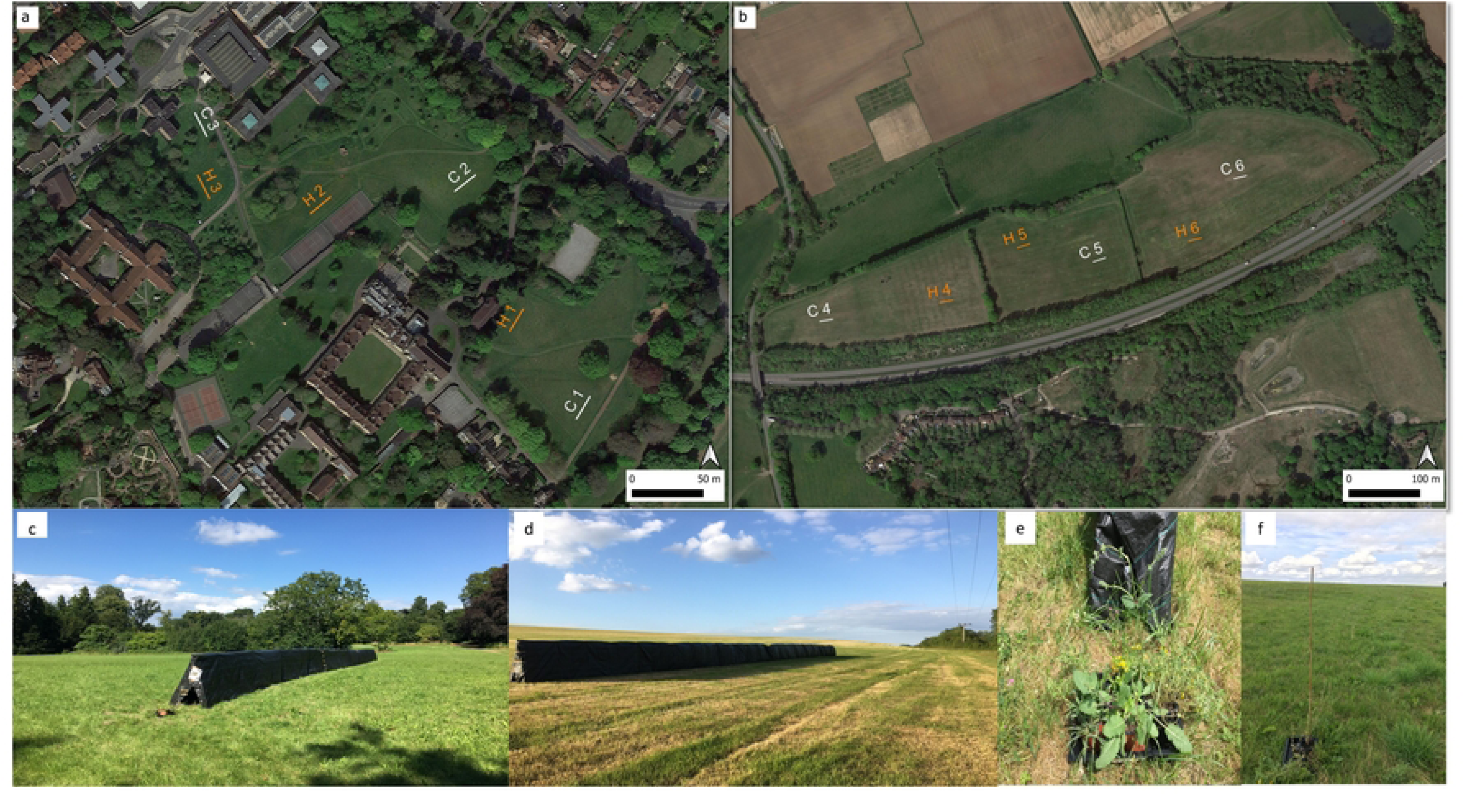
Study site and experimental setup. Aerial photograph shows the study sites in the urban (a) and rural (b) areas. Numbered orange lines show the locations of the artificial linear features and numbered white lines show the controls. Artificial linear features were constructed in both urban sites (c) and rural sites (d). Experimental plant communities were placed at either end of the artificial linear features (e) and controls (f).

### 2.2 Construction of the linear features

Six linear features were constructed in six open patches of meadows at both field sites. The linear features were at least 100m from each other and were considered as independent replicates. Each linear feature was placed in the centre of each meadow space and they were 30 m in length and 1.2 m in height, and made from a bamboo framework covered with black weed suppressing matting [Figure 1c & d, similar to 11]. These linear features provide good visibility compared with other ground-level landmarks, but without providing any floral resources. Our phytometer plant communities (see below) were placed at either end of the linear features. These were then compared to phytometer plants, which were unconnected by linear features, rather each plant community was simply placed 30 m apart. The controls were at least 30m away from the linear features. Halfway through the field season, we swapped the positions of the linear features and the controls (i.e., moved them to the where the controls were placed and *vice versa*) to avoid potential any confounding impact of spatial location.

### 2.3 Experimental plant communities

We assembled replicate phytometer communities of seven plant species. Plant species were chosen to be largely pollinated by specific functional groups of pollinators, i.e., bees, flies, and moths. The bee pollinated species were *Salvia verbenaca*, *Lotus corniculatus*, and *Borago officinalis* [35–37], the fly pollinated species were *Centaurium erythraea* and *Anthemis arvensis* [38, 39], and the moth pollinated species were *Silene nutans* and *Zaluzianskya capensis* [40, 41]. The plants were all native to the UK, with the exception of *Z. capensis* which is native to South Africa and was used to provide replication of a moth pollinated plant. There were no plants of these species observed within 100m radius of each site for most species, except for *L. corniculatus*, which was a common species widely distributed in lowland pastures. To mitigate potential confronting effects, we manually removed the flowers of any individuals of this species occurred within 100m radius of the phytometers multiple times during the field season. Plant communities were observed for a limited amount of time to confirm diurnal pollinator assemblages (Fig S1).

Two of the species, *L. corniculatus* and *A. arvensis,* are known to be self-incompatible [39, 42], four species, *S. nutans*, *S. verbenaca*, *B. officinalis*, and *C. erythraea,* are self-compatible to some degree [37, 43–45] and the reproductive system of the remaining species *Z. capensis* was unknown. While self-compatible plants can set seeds in the absence of pollinators, a higher or better seed set will be expected when visited by pollinators [46]. To investigate the extent to which the seed set of each species was affected by open pollination, we conducted a bagging experiment where 5-10 individuals of each species were bagged to exclude pollinator visitations. The seed set of individuals from pollinator exclusion were harvested and counted thereafter. Open pollination is known to improve seed weight of *S. verbenaca* [37], therefore we measured the effect of pollination on the seed set of this species by calculating the average weight per seed.

### 2.4 Using plant communities to measure pollination

Plants were grown in the greenhouse until flowering size and pollinators were excluded during this period. Each species was sowed as the same cohort under uniform lighting and nutrient conditions. Once flowering, they were placed as potted plants in small trays, in a community of seven species in pairs 30m apart, either linked by a linear feature, or not if they were the controls. The trays were used to make watering in the field easier. One individual plant of each species was placed in a tray, thus for every species, a total of 24 individuals were put out in the treatments and controls per round of sampling (6 linear features plus 6 controls, i.e. 12 plants, each with two ends = 24 plants). *S. verbenaca* and *S. nutans* both had an early and short flowering season and were put into the field for one round with 24 individuals of each species, whilst the other species with longer or late flowering season (i.e., *L. corniculatus*, *B. officinalis*, *A. arvensis*, *C. erythraea*, *Z. capensis*) had two or three rounds totalling 48 or 72 individuals over the field season (Fig. S2). Although we could not place all the species outside at the same time due to variation in flowering phenology, there were always at least three species in each plant community during the experiment. Each plant was placed outside until their petals dropped; for the majority of species this happened during the first two weeks, expect*. capensis* which was in the field for three weeks. Flowers of different sizes were randomized between the treatment and control with their flower numbers were counted when placing outside as starting densities. There were no significant differences in the starting flower density of each species between the linear features and control (Table S1). Once flowering was over, the plants were returned to the greenhouse to set seed, and pollinators were excluded to avoid any further pollination occurring. Plants were that dead or damaged were removed from analysis and any flowers which opened after the plants returned to the greenhouses were removed.

As having a single phytometer plant per species in each community could result in low observed pollinator visitation per plant species (Figure S3), we used the seed set of per plant as a measure of pollination. Seeds were harvested as they ripened over the growing season. We counted the total number of seeds collected from each individual plant, and weighted dry biomass. It was impossible to measure the number and average weight of seeds produced by*C. erythraea* and *A. arvensis* as they produce a very large number of very tiny seeds per capsule, therefore only seed biomass was measured for these species.

### 2.5 Data analysis

All statistical analysis were conducted in R [v4.0.2, 47].

#### 2.5.1 Pollinator exclusion

We first investigated the effect of open pollination on the seed set of the five self-compatible species by comparing the seed set of plants from pollination exclusion with same species placed in the field. A non-parametric Manny-Whiteney *U* test was performed for each species, incorporating seed numbers for *B. officinalis, Z. capensis*, *S. nutans,* seed biomass for *C. erythraea*, and seed quality for *S. verbenaca*. The effect of pollinator exclusion was not tested on the two self-incompatible species, *A. arvensis* and *L. corniculatus*.

#### 2.5.2 Species-level response

We used a two-way analysis of variance (ANOVA) to investigate if the presence of linear feature (presence vs absence) and habitat type (i.e., urban vs. rural) affected the seed set of each species. Data were transformed using natural logarithm (except the seed quality of *S. verbenaca* and seed number of *Z. capensis* which was square root transformed) to meet the model assumption. We included linear features, habitat types, and their interaction as predictors. Model residuals were checked using Shapiro-Wilk test and homogeneity of variances was tested using Levene’s test. Tukey’s post hoc test was performed to compare group level differences.

#### 2.5.3 Community-level response

We used non-metric multidimensional scaling (NMDS) in ‘Vegan’ package [48] to investigate the community-level response of seed set to linear feature and habitat type. As *S. verbenaca* and *S. nutans* were placed for one round with 24 individuals each, while other species were placed two or three rounds of 48 or 96 individuals, we standardized seed set measures of each species to 24 individuals by averaging the seed set measures of each species over the rounds. As plants were measured for different seed set traits (i.e., seed numbers, seed biomass, seed quality), we quantified the contribution of pollination to the increase in the seed set trait for each species, through dividing each trait value by the average seed measure collected from the pollinator exclusion experiment:

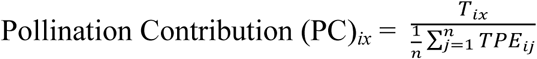

where T *_ix_* is the trait measure for species *i* at community id *x*, and TPE *_ij_* is the trait measured at n number of samples in the pollination exclusion for species *i*. For two self-incompatible species *A. arvensis* and *L. corniculatus* where pollination exclusion was not tested, we used the trait value of each species, divided by the mean trait value calculated from the samples that were placed in the field without linear feature connections. The seed numbers of *Z. capensis* were standardized to 67% as this species was placed outside for three weeks while other species were two weeks (Figure S1). Thus, plants receiving more pollination will set more or heavier seeds relative to pollinator exclusion, which indicates a higher pollination contribution to the seed set trait of this species.

We then calculated the Bray-Curtis dissimilarity matrices for each pair of communities to quantify overall compositional dissimilarity of pollination contribution between any two communities, and compare if the seed set outputs of these plant communities differed due to the landscape effect (i.e., linear features, habitat) on pollination. As pollination contribution was quantified as the increase in seed set traits (e.g., number, biomass, average seed weight) in relative to pollinator exclusion, the dissimilarities matrices indicated the differences in the seed set increase between plant communities. Any communities containing missing values due to loss of samples were removed from the analysis. As there were no evidence that the seed set of *S. nutans* responded to pollination (see results), we excluded this species from the analysis. NMDS was performed at two dimensions of ordination (k = 2) using ‘metaMDS’ function. The association between plant species and ordinations was extracted using ‘envfit()’ function. To provide a broad picture of how the presence of linear features affected the pollination of plant communities at urban and rural site respectively, we grouped samples by their attributes into the following groups urban/artificial linear features, rural/artificial linear features, urban/no linear features, and rural/no linear features. A permutational multivariate analysis of variance (PERMANOVA) was used to compare community-level differences in pollination, incorporating artificial linear features, habitat, and their interaction as predictors. Collectively, the association between plant species and ordination will indicate which species correlate with the largest fraction of pollination contribution within the community, and the PERMANOVA will indicate if the increase in the seed set outputs (relative to pollinator exclusion) of plant communities were different between linear features and control, and between urban and rural. Finally, the NMDS ordination plot will visualize the similarity/dissimilarity of pollination in plant communities.

## 3. Results

### 3.1 Pollinator exclusion

Pollinator exclusion significantly reduced the seed set of four of the five self-compatible species. Specifically, pollinator exclusion reduced 88.6 % of mean seed numbers in bee-pollinated *B. officinalis* (Mann–Whitney *U* test, W = 216, *p* < 0.001), and 17.9% of average seed weight in *S. verbenaca* (Mann–Whitney *U* test, W = 121, *p* = 0.003). For fly- and moth-pollinated species, pollinator exclusion reduced 53.8 % of average seed weight in *C. erythraea* (Mann–Whitney *U* test, W = 232.5, *p* = 0.007) and 81.7% of seed number in *Z. capensis* (Mann–Whitney *U* test, W = 246, *p* = 0.03). However, open pollination did not increase the seed numbers collected from *S. nutans* (Mann–Whitney *U* test, *p* > 0.05). Overall, open pollination increased the seed set of six of the seven species used in the study.

### 3.2 Species-level response

The presence of linear features significantly improved the seed set of the three bee-pollinated species, with higher number of seeds produced in *B. officinalis* and *L. corniculatus*, and larger weight per seed in *S. verbenaca* (Table 1, Table 2). No effect of habitat (urban vs rural) was found in bee pollinated species (Table 1). Linear features did not significantly affect the seed biomass of fly pollinated *C. erythraea* or *A. arvensis,* nor the seed number of moth pollinated *Z. capensis* and *S. nutans* (Table 1). However, the seed biomass of *C. erythraea* and the seed number of *Z. capensis* were significantly affected by habitat (Table 1), with higher seed set found in the urban study site than rural study site (Table 2). Furthermore, pair-wise analysis showed that when connected by linear features, there was higher number of seeds produced by *Z. capensis* in the urban than rural (Tukey’s HSD, *p* = 0.033, Table S3). Finally, there were no significant interactions between habitats and linear feature on the seed set for any species (Table 1).

**Table 1.**
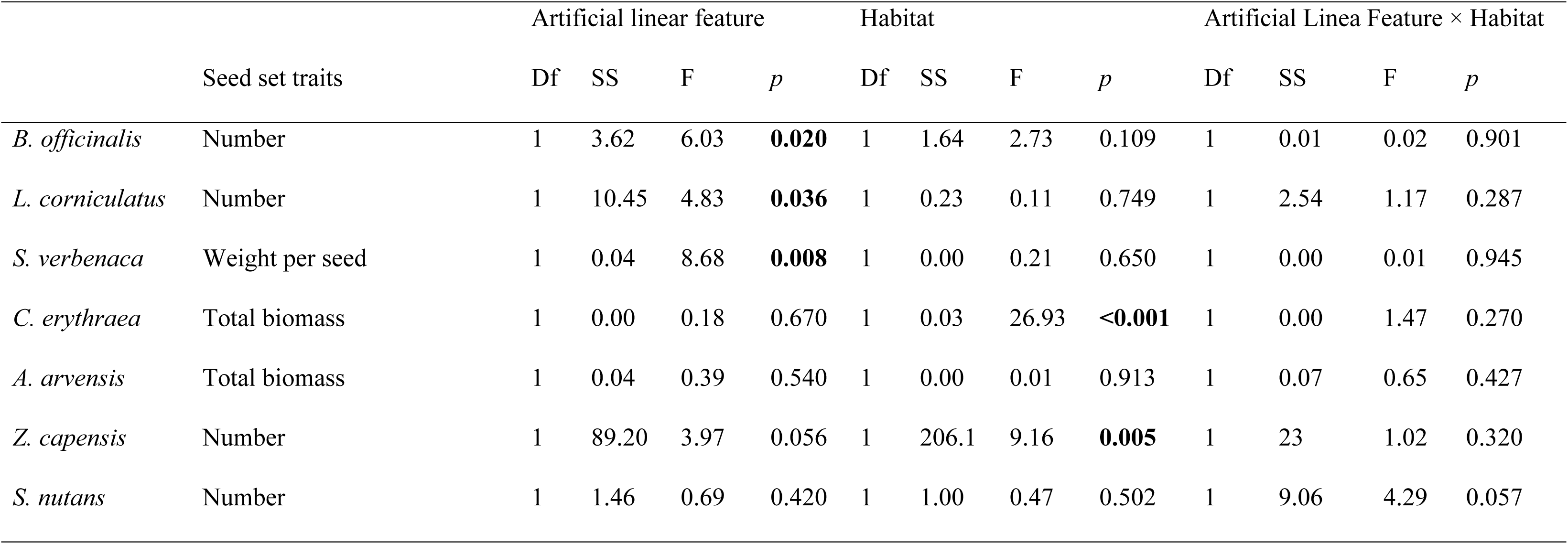
Summary of the two-way ANOVA on the seed set traits of each species. Data were transformed on a log-transformed scale (where average seed weight of *S. verbenaca* and seed number of *Z. capensis* were square-root transformed). Significant effects (*p* < 0.05) were denoted as bold.

**Table 2.**
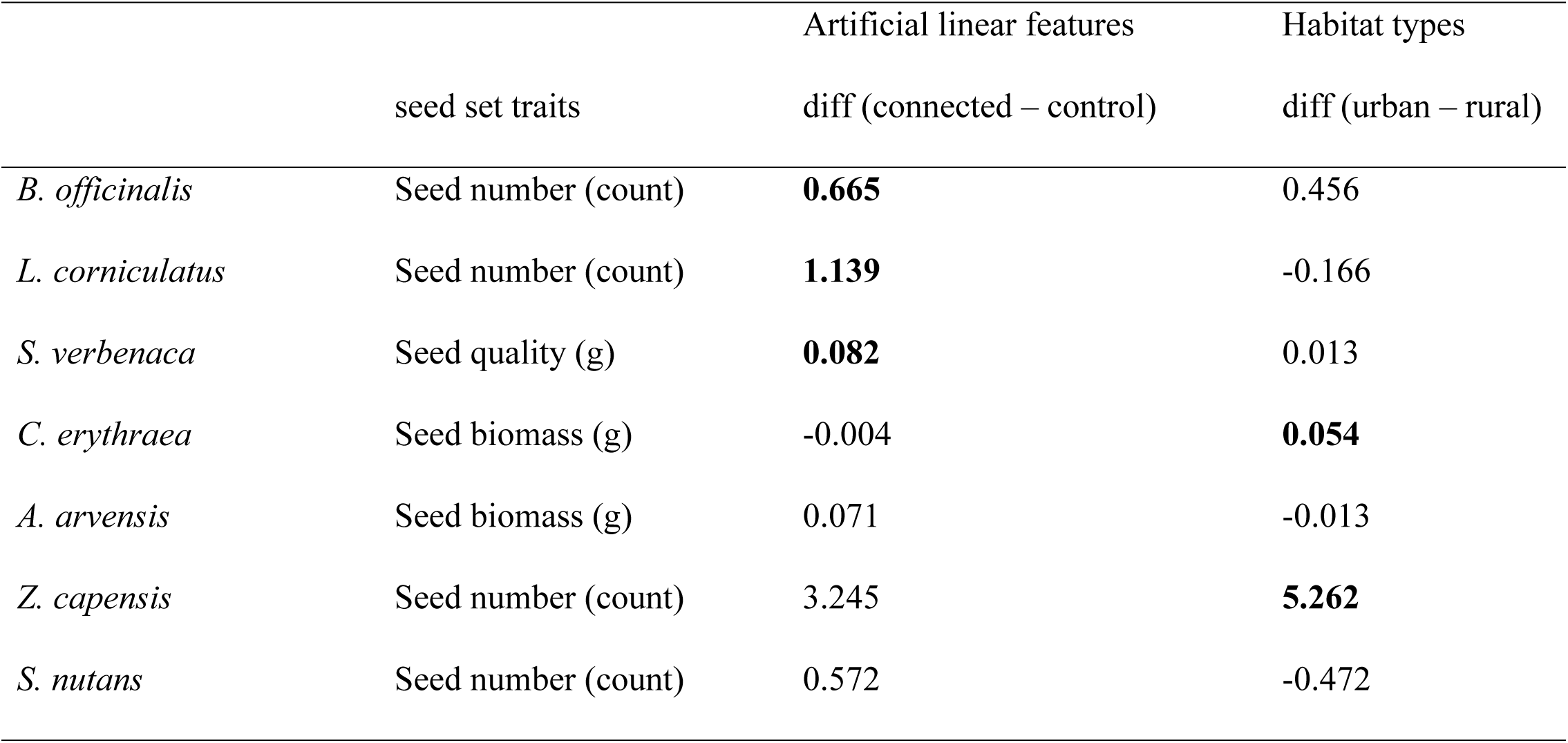
Pairwise differences in the seed set traits of phytometer species. *diff* indicates the differences in group-level mean on a log-transformed scale (where seed biomass of *S. verbenaca* was square-root transformed). Significant differences of mean values (*p* < 0.05) were denoted as bold.

### 3.3 Community-level response

Both artificial linear features and habitat significantly affect the pollination of plant communities (Linear features: F_1,18_ = 2.642, *p* = 0.043; Habitat: F_1,18_ = 3.252, *p* = 0.023). Specifically, connecting plant communities with linear features at urban resulted in different amount of increase in seed set outputs at the community level, compared with other three groups (Fig. 3). *B. officinalis* and *Z. capensis* were strongly associated with the ordination (*B. officinalis*: R^2^ = 0.918, *p* = 0.001; *Z. capensis:* R^2^ = 0.8358, *p* = 0.001), and contributed the largest fractions of pollination contributions within the community (Fig. 2).

**Fig 2.**
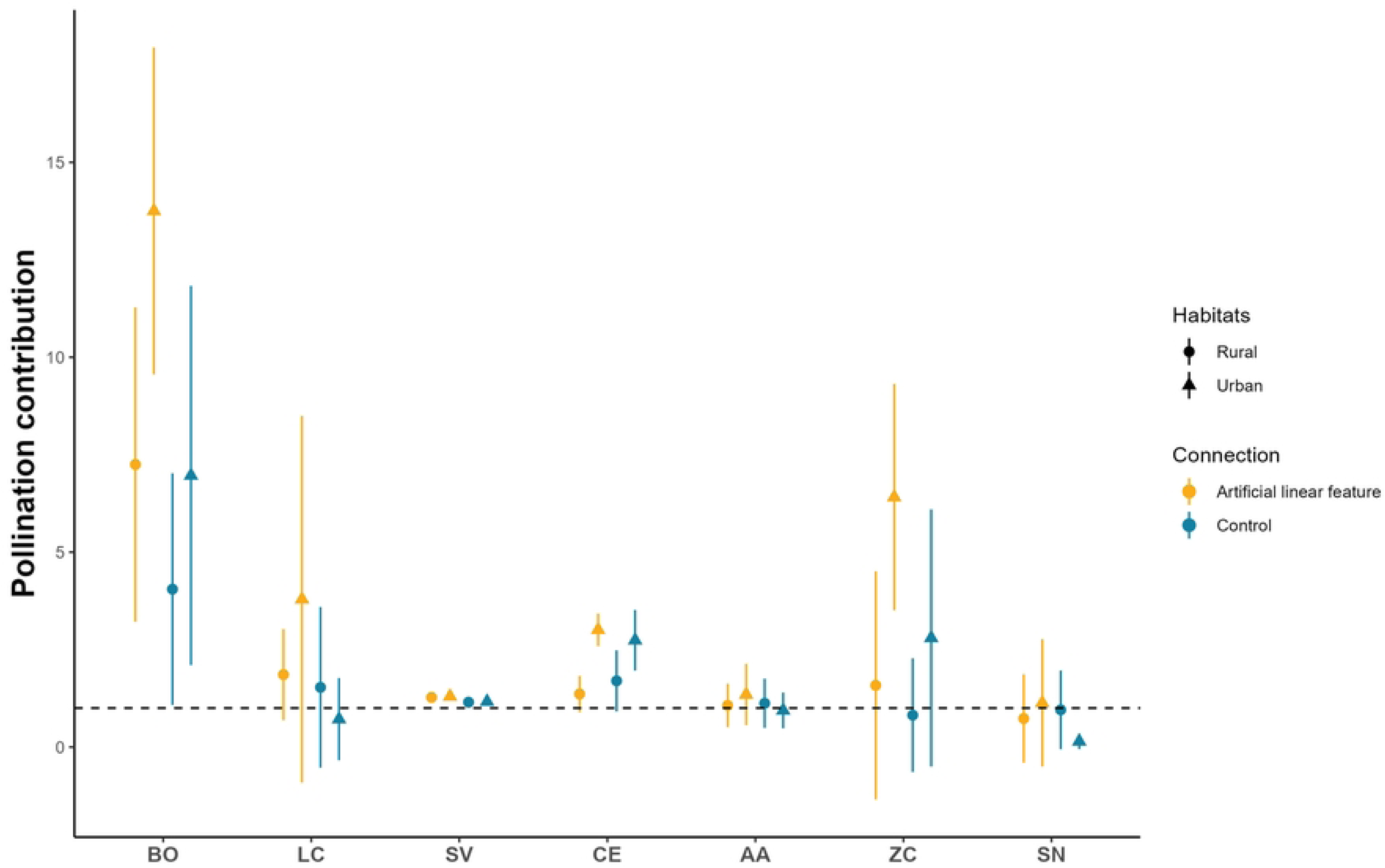
The pollination contribution of seven phytometer species in the plant community. Pollination contribution was calculated in relative to the referenced seed set traits in the pollinator exclusion (dashed line). Plant communities were placed either in urban (triangle) or rural (dot), and either connected by artificial linear features (yellow) or unconnected (blue). Bee-pollinated species: BO = *B. officinalis*; LC = *L. corniculatus*; SV = *S. verbenaca*; Fly-pollinated species: CE = *C. erythraea*; AA = *A. arvensis*; moth-pollinated species: ZC = *Z. capensis*; SN = *S. nutans*.

**Fig 3.**
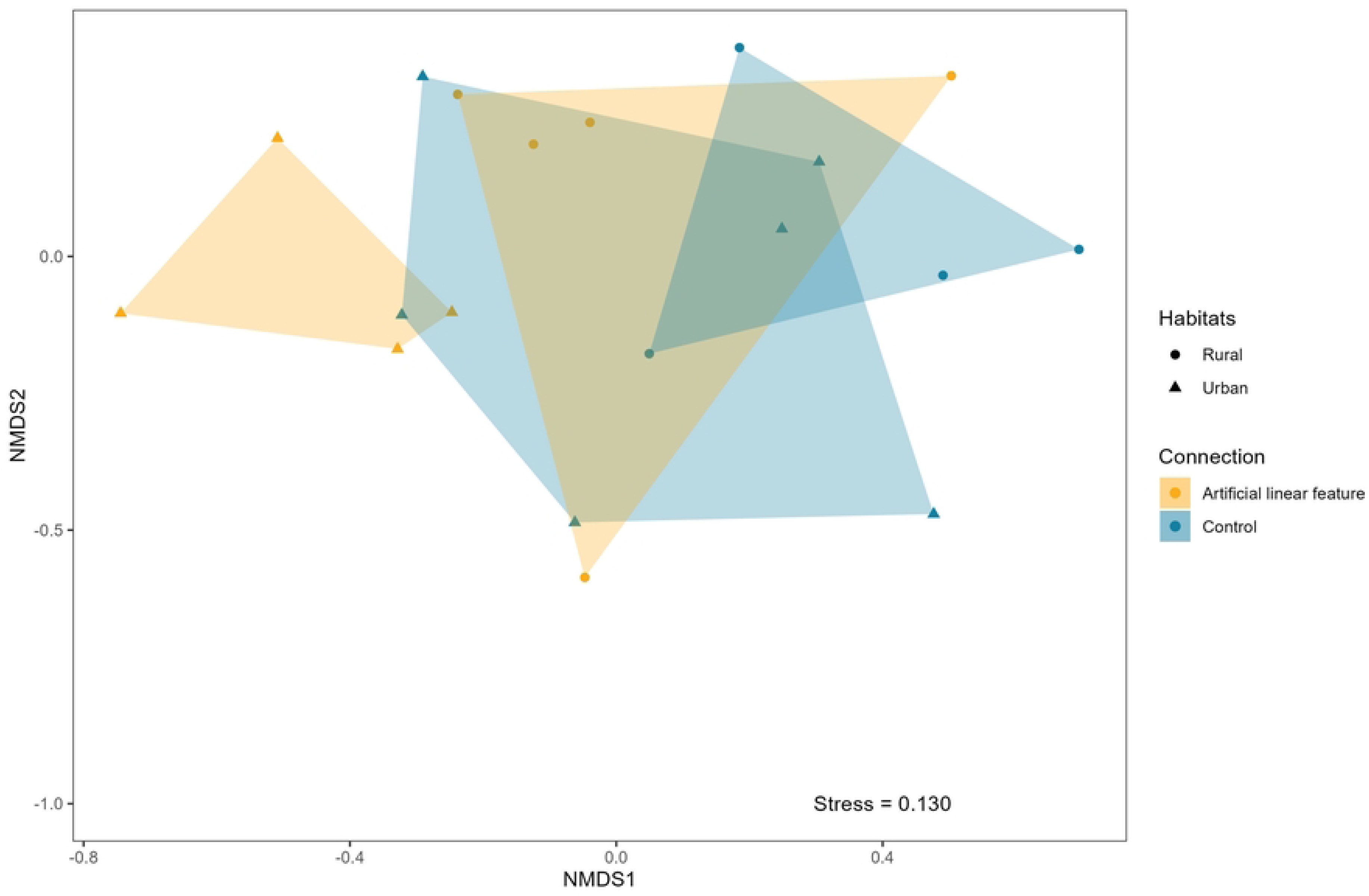
Two-dimensional NMDS ordination plot (stress = 0.130). Showed the clustering the pollination of plant communities either connected with artificial linear features (yellow) or control (blue), and placed in either urban (dots) or rural (triangles) habitats.

## 4. Discussion

Using plant communities consisting of phytometer species that differed in pollination traits, we measured the impact of linear features on pollinator communities and tested if this effect was different between urban and rural habitats. Our results showed that connecting plant communities with linear features could increase the pollination success of bee-pollinated plants, whereas the pollination fly and moth pollinated plants was not significantly improved. This insect order effect did not differ in habitats, though a higher pollination success were found in urban habitats for two of seven species. Furthermore, there were significant differences in community level response of pollination between urban and rural habitats, and between communities connected with a linear feature and not connected. Specifically, we found that connecting communities with linear features at urban habitat resulted in different amount of increase in seed set outputs at the community level, relative to the amount of seed set produced when pollinators were excluded. In which follows, we discuss the potential limitations to our approach, and the impact of linear features on the pollination of different pollinator groups, highlighting the importance of using community-level approach to study the impact of landscape features on pollinator communities.

### 4.1 Limitations to approach

The purpose of this study was to use a logistically feasible approach to test the role of linear features in pollination with replicate plant communities. While we approached this with suitable levels of replication, each type of study site (urban vs rural) was only replicated once, which prevent us from drawing broad conclusions on how different habitats affected pollinator assemblages along with the presence of linear features. Although it is unlikely that pollinator compositions and richness at one certain habitat were vastly different across years, rather varying in different landscape scales, we acknowledged that replicating at a broader geographical range or across years may provide more comprehensive evidence. This also stresses the needs for evaluating other pollinator taxa such as beetles, ants, and wasps, though their importance in biotic pollination is recognised [49, 50]. While the pollination by these groups is relatively rare [51], and specialist plants pollinated by these groups are more difficult to find in comparison to bee, fly and moth pollinated plants [52].

### 4. 2 The impact of the linear features on bee, fly and moth pollinated plants

We showed that connecting plant communities with linear features increased the pollination success of all three bee pollinated plants, providing evidence that they functioned as dispersal corridors for bees. Although artificial linear features did not provide any floral resources or habitats, it is possible that they could provide unique visual clues, which may play an important role for bees during foraging [15, 53]. Indeed, bees are central place foragers and need to return their hives once foraging is complete, meaning that finding distinct landscape features would make easier for them to locate patches of floral resources and navigate back to their hive [15, 54] The nature of central place foraging indicated that bees are likely to rely on unique landscape features during dispersal. For example, by tracking pollen movements of primrose *Primula vulgaris* mainly pollinated by bees, Van Geert, Van Rossum [9] found higher rates of pollen dispersal when plants were connected by ditches. This aligns with our findings that the seed set of three phytometer species that were pollinated by bees, increased when plants were connected, this suggesting that linear features could function as corridors the plants they pollinate, as well as the bees themselves.

The magnitude of a corridor effect can vary according to corridor size and quality [3, 9, 55]. In our study here, we placed plant communities 30m as that fitted our field sites and seemed a reasonable place to start. The effect of linear feature lengths on the movement of bee pollinators was beyond the scope of this study, though an area worthy of further investigation and likely predicted in part by the dispersal ability of the these pollinator groups [56, 57].

We found that connecting our phytometer communities with an artificial linear feature did not affect the seed production of our two fly pollinated species. Although travelling 30m distance is probably not difficult for most flower-visiting flies, there is no evidence that flies were able to use these features to disperse. There is relatively little evidence on the dispersal patterns of flies [but see 23], probably because flies are highly dispersive and difficult to observe. In fact, the movement of majority fly pollinators, such as hoverflies, are not central place foragers and often seen as ‘one-way’ movement during foraging [58, 59]. This suggests that some linear habitats with substantial resources, such as hedgerows [60, 61] and road verges [10], are likely to host more Diptera pollinators, may because they contain forage. In our study, the artificial linear feature provided no resources, and unlike bees, following linear features without floral rewards is unlikely to provide many benefits for flies [62].

In contrast to previous findings that linear landscape features functioned as dispersal corridors for moth pollinators [12, 25, 63], we found that although the linear features did not affect the seed set of the moth pollinated species *Z*. *capensis*, connecting plant communities with a corridor increased its seed set in the urban, but not the rural site. However, as open pollination did significantly not increase seed set compared with pollinator exclusion, it is difficult to compare the effect of the linear features on the other moth pollinated plant*, S. nutans*. It could be due to that *S. nutans* was not pollinated as this species was only replicated in the field for one round of placement (Fig. S1). This means that if moth activities were low in the site during certain time of the year, plants may not be visited enough to gain effects on seed set. However, the positive effect of habitats on the seed set of *Z*. *capensis* when communities were connected suggested that the effect of artificial linear features on moth pollinators may be site specific. Given the contrasting results provided by two moth pollinated species, more research is needed to be completed before any conclusions can be reached.

### 4. 3 The impact of artificial linear features at the community level

The differing responses of pollinator groups and plant species to the linear features highlights the importance of measuring a community-wide response to the feature, rather than focusing on its impact on a single plant species (usually one visited by bees). While this focus can provide useful information, plant-pollinator communities consist of many interacting species and investigating just species-level dynamics may overlook the impacts of corridors on pollination [28]. By using communities of plant species with complimentary pollinator traits, we quantified the impact of artificial linear features on the pollination of a plant communities to evaluate their roles as biological corridors for pollinator communities. We found that connecting plant communities with artificial linear features facilitated the movements of bees and increased bee pollination of three species, whilst fly and moth pollinators remained unaffected or less affected. Moreover, the differences in pollination contribution within plant communities indicated that some plants might benefit more than other species. These results demonstrate that the different responses of pollinators to the same landscape structure could affect plant community dynamics and their response to habitat change. Damschen, Brudvig [64] reported that plant seed dispersal was affected by the responses of seed carriers to corridors, and likewise, our study highlights that plant pollination is affected by the response of the pollinators to corridors. It is worth noting that many plant species require *both* biotic pollination and biotic seed dispersal, and so the situation is in fact more complex than shown by either Damschen et al. (2008) or by our project reported here.

### 4.4 Implications for conservation

Our results have implications for the conservation of insect pollinators and their pollination. Bees can use linear features as flight paths for navigating between the nest and foraging areas and other ground-level linear elements such as hedges, ditches, walls are likely to act as movement conduits for them. Thus, conserving linear landscape features is likely to be useful for the plants they pollinate and the bees themselves. Diptera and Lepidoptera are important pollinators [e.g., 65, 66] but they do not appear to respond to linear features and other conservation measures may prove more useful for them, for example stepping stones of foraging or breeding habitats. That said, more studies are needed to evaluate how the matrix between focal habitats affects Diptera and Lepidoptera.

## 5. Conclusions

While linear features are very likely to improve pollination for some groups of species, our results suggest that this effect is inconsistent, thus we found beneficial impacts for bee pollinated plants but not for fly or moth pollinated plants. By studying pollination success in what are effectively small model communities, it is possible to gain some insights about how ecological processes such as pollination are likely be affected by habitat fragmentation at the plant community level.

## Acknowledgement

We thank Alanna Kelly and Tom Pitman for help with growing and maintaining the plants used in the experiment. This work was supported by the China Scholarship Council (Grant no. 20186190011).

## Data availability statement

Data are available on GitHub repository (https://github.com/Dongboli/experimental-data) for peer review but will make the data available once the paper is published, using figshare repository.

## Supporting information

S1 file. Supporting information of the impact of artificial linear features on the pollination of plant communities.

## References

1. Lindenmayer DB, Fischer J. Tackling the habitat fragmentation panchreston. Trends in Ecology & Evolution. 2007;22(3):127–32. doi: 10.1016/j.tree.2006.11.006.

2. Haddad NM, Brudvig LA, Clobert J, Davies KF, Gonzalez A, Holt RD, et al. Habitat fragmentation and its lasting impact on Earth’s ecosystems. Science advances. 2015;1(2):e1500052. doi: 10.1126/sciadv.150005.

3. Li D, Clements CF, Shan IL, Memmott J. Corridor quality affects net movement, size of dispersers, and population growth in experimental microcosms. Oecologia. 2021;195:547–56. doi: 10.1007/s00442-020-04834-2.

4. Christie MR, Knowles LL. Habitat corridors facilitate genetic resilience irrespective of species dispersal abilities or population sizes. Evolutionary Applications. 2015;8(5):454–63. doi: 10.1111/eva.12255.

5. Henein K, Merriam G. The elements of connectivity where corridor quality is variable. Landscape ecology. 1990;4:157–70. doi: 10.1007/BF00132858.

6. Habel JC, Ulrich W, Schmitt T. Butterflies in corridors: quality matters for specialists. Insect Conservation and Diversity. 2020;13(1):91–8. doi: 10.1111/icad.12386.

7. Resasco J, Haddad NM, Orrock JL, Shoemaker D, Brudvig LA, Damschen EI, et al. Landscape corridors can increase invasion by an exotic species and reduce diversity of native species. Ecology. 2014;95(8):2033–9. doi: 10.1890/14-0169.1.

8. Söderström B, Hedblom M. Comparing movement of four butterfly species in experimental grassland strips. Journal of insect Conservation. 2007;11:333–42. doi: 10.1007/s10841-006-9046-5.

9. Van Geert A, Van Rossum F, Triest L. Do linear landscape elements in farmland act as biological corridors for pollen dispersal? Journal of Ecology. 2010;98(1):178–87. doi: 10.1111/j.1365-2745.2009.01600.x.

10. Dániel-Ferreira J, Berggren Å, Wissman J, Öckinger E. Road verges are corridors and roads barriers for the movement of flower - visiting insects. Ecography. 2022;2022(2). doi: 10.1111/ecog.05847.

11. Cranmer L, McCollin D, Ollerton J. Landscape structure influences pollinator movements and directly affects plant reproductive success. Oikos. 2012;121(4):562–8. doi: 10.1111/j.1600-0706.2011.19704.x.

12. Coulthard E, McCollin D, Littlemore J. The use of hedgerows as flight paths by moths in intensive farmland landscapes. Journal of Insect Conservation. 2016;20:345–50. doi: 10.1007/s10841-016-9864-z.

13. Dondina O, Kataoka L, Orioli V, Bani L. How to manage hedgerows as effective ecological corridors for mammals: a two-species approach. Agriculture, Ecosystems & Environment. 2016;231:283–90. doi: 10.1016/j.agee.2016.07.005.

14. Pelletier-Guittier C, Théau J, Dupras J. Use of hedgerows by mammals in an intensive agricultural landscape. Agriculture, Ecosystems & Environment. 2020;302:107079. doi: 10.1016/j.agee.2020.107079.

15. Brebner, Makinson JC, Bates OK, Rossi N, Lim KS, Dubois T, et al. Bumble bees strategically use ground level linear features in navigation. Animal Behaviour. 2021;179:147–60. doi: 10.1016/j.anbehav.2021.07.003.

16. Van Rossum F, Triest L. Stepping-stone populations in linear landscape elements increase pollen dispersal between urban forest fragments. Plant Ecology and Evolution. 2012;145(3):332–40. doi: 10.5091/plecevo.2012.737.

17. Anderson M, Rotheray EL, Mathews F. Marvellous moths! pollen deposition rate of bramble (Rubus futicosus L. agg.) is greater at night than day. Plos one. 2023;18(3):e0281810. doi: 10.1371/journal.pone.0281810.

18. Walton RE, Sayer CD, Bennion H, Axmacher JC. Nocturnal pollinators strongly contribute to pollen transport of wild flowers in an agricultural landscape. Biology letters. 2020;16(5):20190877. doi: 10.1098/rsbl.2019.0877.

19. Rader R, Bartomeus I, Garibaldi LA, Garratt MP, Howlett BG, Winfree R, et al. Non-bee insects are important contributors to global crop pollination. Proceedings of the National Academy of Sciences. 2016;113(1):146–51. doi: 10.1073/pnas.1517092112.

20. Klein S, Cabirol A, Devaud J-M, Barron AB, Lihoreau M. Why bees are so vulnerable to environmental stressors. Trends in ecology & evolution. 2017;32(4):268–78. doi: 10.1016/j.tree.2016.12.009.

21. Hanski I, Saastamoinen M, Ovaskainen O. Dispersal-related life-history trade-offs in a butterfly metapopulation. Journal of animal Ecology. 2006;75(1):91–100. doi: 10.1111/j.1365-2656.2005.01024.x.

22. Rodríguez-Gasol N, Avilla J, Alegre S, Alins G. Sphaerophoria rueppelli adults change their foraging behavior after mating but maintain the same preferences to flower traits. BioControl. 2019;64(2):149–58. doi: 10.1007/s10526-019-09928-2.

23. Fried JH, Levey DJ, Hogsette JA. Habitat corridors function as both drift fences and movement conduits for dispersing flies. Oecologia. 2005;143:645–51. doi: 10.1007/s00442-005-0023-6.

24. Haddad NM, Tewksbury JJ. Low-quality habitat corridors as movement conduits for two butterfly species. Ecological Applications. 2005;15(1):250–7. doi: 10.1890/03-5327.

25. Várkonyi G, Kuussaari M, Lappalainen H. Use of forest corridors by boreal Xestia moths. Oecologia. 2003;137:466–74. doi: 10.1007/s00442-003-1354-9.

26. Öckinger E, Smith HG. Do corridors promote dispersal in grassland butterflies and other insects? Landscape ecology. 2008;23:27–40. doi: 10.1007/s10980-007-9167-6.

27. Dover J, Fry G. Experimental simulation of some visual and physical components of a hedge and the effects on butterfly behaviour in an agricultural landscape. Entomologia experimentalis et applicata. 2001;100(2):221–33. doi: 10.1046/j.1570-7458.2001.00867.x.

28. Willcox BK, Aizen MA, Cunningham SA, Mayfield MM, Rader R. Deconstructing pollinator community effectiveness. Current Opinion in Insect Science. 2017;21:98–104. doi: 10.1016/j.cois.2017.05.012.

29. Hackett TD, Sauve AM, Maia KP, Montoya D, Davies N, Archer R, et al. Multi-habitat landscapes are more diverse and stable with improved function. Nature. 2024;633(8028):114-9. doi: 10.1038/s41586-024-07825-y.

30. Orford KA, Murray PJ, Vaughan IP, Memmott J. Modest enhancements to conventional grassland diversity improve the provision of pollination services. Journal of Applied Ecology. 2016;53(3):906–15. doi: 10.1111/1365-2664.12608.

31. Li D, Clements CF, Memmott J. Isolation limits spring pollination in a UK fragmented landscape. Plos one. 2024;19(9):e0310679. doi: 10.1371/journal.pone.0310679.

32. Dietzel S, Rojas-Botero S, Kollmann J, Fischer C. Enhanced urban roadside vegetation increases pollinator abundance whereas landscape characteristics drive pollination. Ecological Indicators. 2023;147:109980. doi: 10.1016/j.ecolind.2023.109980.

33. Fontaine C, Dajoz I, Meriguet J, Loreau M. Functional diversity of plant–pollinator interaction webs enhances the persistence of plant communities. PLoS biology. 2006;4(1):e1. doi: 10.1371/journal.pbio.0040001.

34. Young CH, Jarvis PJ. Measuring urban habitat fragmentation: an example from the Black Country, UK. Landscape Ecology. 2001;16:643–58. doi: 10.1023/A:1013108005347.

35. Descamps C, Quinet M, Baijot A, Jacquemart AL. Temperature and water stress affect plant–pollinator interactions in Borago officinalis (Boraginaceae). Ecology and evolution. 2018;8(6):3443–56. doi: 10.1002/ece3.3914.

36. Jones DA, Turkington R. Biological flora of the British Isles - Lotus corniculatus L. Lotus corniculatus. 1986:1185–212. doi: http://www.jstor.org/stable/2260243.

37. Navarro L. Is the dichogamy of Salvia verbenaca (Lamiaceae) an effective barrier to self-fertilization? Plant Systematics and Evolution. 1997;207:111–7. doi: 10.1007/BF00985212.

38. Brys R, Jacquemyn H. Variation in the functioning of autonomous self-pollination, pollinator services and floral traits in three Centaurium species. Annals of Botany. 2011;107(6):917–25. doi: 10.1093/aob/mcr032.

39. Kay Q. Biological flora of British Isles - Anthemis arvensis L. Journal of Ecology. 1971;59(2):637-&. doi: 10.2307/2258337.

40. Johnson S, Edwards T, Carbutt C, Potgieter C. Specialization for hawkmoth and long- proboscid fly pollination in Zaluzianskya section Nycterinia (Scrophulariaceae). Botanical Journal of the Linnean Society. 2002;138(1):17–27. doi: 10.1046/j.1095-8339.2002.00005.x.

41. Vanderplanck M, Touzet P, Van Rossum F, Lahiani E, De Cauwer I, Dufaÿ M. Does pollination syndrome reflect pollinator efficiency in Silene nutans? Acta Oecologica. 2020;105:103557. doi: 10.1016/j.actao.2020.103557.

42. Pellissier V, Muratet A, Verfaillie F, Machon N. Pollination success of Lotus corniculatus (L.) in an urban context. Acta Oecologica. 2012;39:94–100. doi: 10.1016/j.actao.2012.01.008.

43. Brys R, De Crop E, Hoffmann M, Jacquemyn H. Importance of autonomous selfing is inversely related to population size and pollinator availability in a monocarpic plant. American Journal of Botany. 2011;98(11):1834–40. doi: 10.3732/ajb.1100154.

44. Lahiani E, Touzet P, Billard E, Dufay M. When is it worth being a self- compatible hermaphrodite? Context - dependent effects of self - pollination on female advantage in gynodioecious Silene nutans. Ecology and Evolution. 2015;5(9):1854–62. doi: 10.1002/ece3.1410.

45. Montaner C, Floris E, Alvarez J. Is self-compatibility the main breeding system in borage (Borago officinalis L.)? Theoretical and Applied Genetics. 2000;101:185–9. doi: 10.1007/s001220051467.

46. Aizen MA, Ashworth L, Galetto L. Reproductive success in fragmented habitats: do compatibility systems and pollination specialization matter? Journal of Vegetation Science. 2002;13(6):885–92. doi: 10.1111/j.1654-1103.2002.tb02118.x.

47. RCoreTeam. R: A language and environment for statistical computing. R Foundation for Statistical Computing, Vienna, Austria. (https://www.R-project.org/). 2022.

48. Oksanen J, Blanchet FG, Kindt R, Legendre P, Minchin PR, O’hara R, et al. Package ‘vegan’. Community ecology package, version. 2013;2(9):1–295. doi: https://github.com/vegandevs/vegan.

49. Rader R, Cunningham S, Howlett BG, Inouye D. Non-bee insects as visitors and pollinators of crops: Biology, ecology, and management. Annual review of entomology. 2020;65(1):391–407. doi: 10.1146/annurev-ento-011019-025055.

50. Requier F, Pérez-Méndez N, Andersson GK, Blareau E, Merle I, Garibaldi LA. Bee and non-bee pollinator importance for local food security. Trends in ecology & evolution. 2023;38(2):196–205. doi: 10.1016/j.tree.2022.10.006.

51. Dutton EM, Frederickson ME. Why ant pollination is rare: new evidence and implications of the antibiotic hypothesis. Arthropod-Plant Interactions. 2012;6(4):561–9. doi: 10.1007/s11829-012-9201-8.

52. Bernhardt P. Convergent evolution and adaptive radiation of beetle-pollinated angiosperms. Pollen and pollination. 2000:293–320. doi: 10.1007/978-3-7091-6306-1_16.

53. Menzel R, Tison L, Fischer-Nakai J, Cheeseman J, Balbuena MS, Chen X, et al. Guidance of navigating honeybees by learned elongated ground structures. Frontiers in Behavioral Neuroscience. 2019;12:322. doi: 10.3389/fnbeh.2018.00322.

54. Collett TS, Graham P. Insect navigation: do honeybees learn to follow highways? Current Biology. 2015;25(6):R240–R2. doi: 10.1016/j.cub.2014.11.003.

55. Haddad N. Corridor length and patch colonization by a butterfly, Junonia coenia. Conservation Biology. 2000;14(3):738–45. doi: 10.1046/j.1523-1739.2000.99041.x.

56. Gathmann A, Tscharntke T. Foraging ranges of solitary bees. Journal of animal ecology. 2002;71(5):757–64. doi: 10.1046/j.1365-2656.2002.00641.x.

57. Greenleaf SS, Williams NM, Winfree R, Kremen C. Bee foraging ranges and their relationship to body size. Oecologia. 2007;153:589–96. doi: 10.1007/s00442-007-0752-9.

58. Wratten, Bowie, Hickman JM, Evans AM, Sedcole JR, Tylianakis JM. Field boundaries as barriers to movement of hover flies (Diptera: Syrphidae) in cultivated land. Oecologia. 2003;134:605–11. doi: 10.1007/s00442-002-1128-9.

59. Meyer B, Jauker F, Steffan-Dewenter I. Contrasting resource-dependent responses of hoverfly richness and density to landscape structure. Basic and Applied Ecology. 2009;10(2):178–86. doi: 10.1016/j.baae.2008.01.001.

60. Garratt MP, Senapathi D, Coston DJ, Mortimer SR, Potts SG. The benefits of hedgerows for pollinators and natural enemies depends on hedge quality and landscape context. Agriculture, Ecosystems & Environment. 2017;247:363–70. doi: 10.1016/j.agee.2017.06.048.

61. Rodríguez-Gasol N, Alins G, Veronesi ER, Wratten S. The ecology of predatory hoverflies as ecosystem-service providers in agricultural systems. Biological Control. 2020;151:104405. doi: 10.1016/j.biocontrol.2020.104405.

62. Jauker F, Diekötter T, Schwarzbach F, Wolters V. Pollinator dispersal in an agricultural matrix: opposing responses of wild bees and hoverflies to landscape structure and distance from main habitat. Landscape Ecology. 2009;24:547–55. doi: 10.1007/s10980-009-9331-2.

63. Balbi M, Croci S, Petit EJ, Butet A, Georges R, Madec L, et al. Least-cost path analysis for urban greenways planning: A test with moths and birds across two habitats and two cities. Journal of Applied Ecology. 2021;58(3):632–43. doi: 10.1111/1365-2664.13800.

64. Damschen EI, Brudvig LA, Haddad NM, Levey DJ, Orrock JL, Tewksbury JJ. The movement ecology and dynamics of plant communities in fragmented landscapes. Proceedings of the National Academy of Sciences. 2008;105(49):19078–83. doi: 10.1073/pnas.0802037105.

65. Macgregor CJ, Pocock MJ, Fox R, Evans DM. Pollination by nocturnal L epidoptera, and the effects of light pollution: a review. Ecological entomology. 2015;40(3):187–98. doi: 10.1111/een.12174.

66. Orford KA, Vaughan IP, Memmott J. The forgotten flies: the importance of non-syrphid Diptera as pollinators. Proceedings of the royal society B: biological sciences. 2015;282(1805):20142934. doi: 10.1098/rspb.2014.2934.

